# A stacked ensemble method for forecasting influenza-like illness visit volumes at emergency departments

**DOI:** 10.1101/2020.10.21.348417

**Authors:** Arthur Novaes de Amorim, Rob Deardon, Vineet Saini

## Abstract

Accurate and reliable short-term forecasts of influenza-like illness (ILI) visit volumes at the emergency departments can improve staffing and resource allocation decisions in each hospital. In this paper, we developed a stacked ensemble model that averages the predictions from various competing methodologies in the current frontier for ILI-related forecasts. We also constructed a back-of-the-envelope prediction interval for the stacked ensemble, which provides a conservative characterization of the uncertainty in the stacked ensemble predictions. We assessed the reliability and accuracy of our model’s 1 to 4 weeks ahead forecasts using real-time hospital-level data on weekly ILI visit volumes during the 2012-2018 flu seasons in the Alberta Children’s Hospital, located in Calgary, Alberta, Canada. Over this time period, our model’s prediction deviated from the realized ILI visit volume by an average of 12% for 1 week ahead forecasts, with a 90% prediction interval having coverage rates ranging from 90.7 to 97.7%.

## Introduction

Influenza-like illness (ILI) causes significant burden to healthcare systems [1, 2]. To address this burden, a growing body of literature focuses on developing accurate and reliable forecasts of ILI to help inform public health decisions and resource planning [3–6]. These research efforts led to multiple candidate models which may be suitable for forecasting ILI activity within a country or region.

While forecasts at a regional level may improve public health decisions, substantial heterogeneity within regions imply a one size fits all forecast at such a large level of spatial aggregation may be of little use for hospital-level staffing and resource allocation decisions [7]. Here, we bridge this literature gap by assessing the forecasting performance of various models from the recent literature using Emergency Department (ED)-level data from the Alberta Real-Time Syndromic Surveillance Net (ARTSSN). Our goal is to equip decision makers in hospitals with near-term forecasts (1 to 4 weeks ahead) of the number of weekly ILI visits to the ED.

We used an ensembling method to leverage the strength of different modeling approaches. Evidence from weather forecasting research [8] and infectious disease forecasts [9] suggest that predictions from ensembling are at least as good as the best performing individual models that they build upon, making the ensemble a potential candidate for predicting ILI positive visit volumes at EDs. We used stacking – a data-driven approach – when leveraging the contribution of each model to the ensemble. In our retrospective analysis, we found that the stacked ensemble model performed as well as – when not better than – the best individual models using various quantitative performance measures on each of the 1 to 4 weeks ahead forecast targets.

## Methods

We performed a retrospective analysis with the objective of forecasting weekly ILI positive visits within an ED. The data and modelling decisions are described below.

### Data

We used data from the ARTSSN, which offers daily records of patients screened ILI positive in Calgary, Alberta. Our sample featured two EDs: the Alberta Children’s Hospital (ACH) – with a high volume of ILI visits – and the Foothills Medical Center (FMC) – with a low volume of ILI visits. The data covered flu seasons from 2012 to 2018, with each flu season starting roughly on the last week of August and lasting 52 or 53 weeks depending on leap year status. The ARTSSN records additionally include patient characteristics such as age, sex, and postal code of last known residence. We supplemented ARTSSN with the following data sets: (1) patients’ recent flu immunization status and dates, obtained from Alberta Health; (2) laboratory diagnostics for ILI-causing viral pathogen confirmation, obtained from the Alberta Public Health Laboratory^1^; (3) annual mid-year population estimates in Calgary, obtained from Calgary’s Civic Census; and, (4) weather information, obtained from Environment and Climate Change Canada.

The unit of observation and analysis in the final linked data set was the weekly-aggregated count of ILI visits at the high-volume ED. We used the low-volume ED data (i.e. FMC) for sensitivity analysis of how the magnitude of visits affects model performance. In some richer model specifications, we further disaggregated the ILI positive visit counts within age brackets and distance buffers around the ED and assessed the impact of these alternative models on forecast performance. Predictors available in a daily format (e.g. minimum temperature) were transformed into weekly averages for the analysis.

### Statistical models

Our objective was to estimate at week *t*, a function 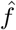 such that the forecast of ILI *ℓ* weeks into the future is given by:

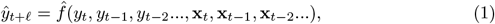

where *y_t_* is the observed count of ILI visits to the ED at week *t* and **x**_*t*_ are other predictors of ILI which are available at time t. Each model drew information from different predictors and specified different relationships between predictors and the outcome of interest. Model implementation is described below.

### Empirical Bayes

The Empirical Bayes model is a framework developed by Brooks *et al* for predicting epidemics by relying on slightly modified versions of past epidemics to form possibilities for the current season [4,10]. With this approach, we first modelled previous flu seasons using non-parametric smoothed piecewise quadratic curves generated with the *cv.trendfilter* method of the *genlasso* R package [11]. This gave us a set of smoothed trajectories {*f^s^*} for each flu season *s* in the training data. Next, we drew at random with equal probability one trajectory from {*f^s^*} and applied a series of transformations to this curve, resulting in a curve 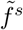. The transformations shift the epidemic curve’s peak height, peak location, and pacing toward the peak, respectively. For these transformations, we randomly sampled a peak height, peak location, and pacing parameter, where candidate peak heights and locations came from the smoothed trajectories *f^s^* and the pacing parameters came from a uniform distribution *U*[0.75, 1.25]. We then assigned a likelihood weight to the transformed curve based on how closely the curve approximated observed ILI up to the current week of the season. We lastly injected noise to the transformed curve, where the noise terms were drawn from a normal distribution 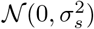 with *σ_s_* being a noise level derived from the trajectory *f^s^*.

The Empirical Bayes prediction of ILI visits ℓ weeks from week *t*, 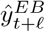, was the weighted median of *N* = 10^5^ random samples of transformed trajectories, with the likelihood weights *w* described above.^2^ Specifically, we ranked the forecast ILI at period *t* + ℓ from the *N* samples from smallest to largest, 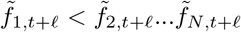 and computed:

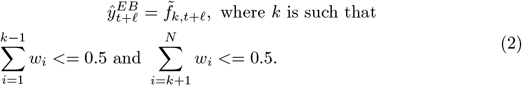

### Autoregressive Integrated Moving Average (ARIMA)

The ARIMA model estimates future periods’ ILI positive visit counts as a function of previous observations and forecast errors. An ARIMA(*p, d, q*) model is specified by three parameters: an autoregressive order term *p*; a degree of differencing *d* for making the time series stationary; and, the order of the moving average *q*. We used seasonal and trend decomposition using locally estimated scatterplot smoothing (STL) to remove seasonality from the raw weekly ILI counts before fitting an ARIMA(2,0,1) model. The analysis was implemented via the *stlm* method of the *forecast* R package [12]. Determination of the number of time lags *p* = 2, degree of differencing *d* = 0, and order of moving average *q* = 1 was based on optimization of the Akaike Information Criterion (AIC) [13].

Upon seasonally adjusting the ILI data with STL, the ARIMA(2,0,1) prediction of ILI positive visits ℓ weeks from week *t*, 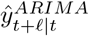, is given by the forecasting equation:

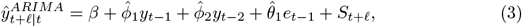

where 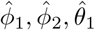 are the estimated model parameters and *S_*t*+ℓ_* is the seasonal component which was computed using STL and removed before fitting the model. The prediction interval is given by 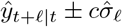 where 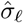 is an estimate of the standard error of the ℓ weeks ahead prediction distribution and *c* comes from the interval coverage probability assuming a normal distribution of forecast errors. Note that the prediction intervals are estimated from the seasonally-adjusted data but may be too narrow as they ignore uncertainty associated with the STL estimation of the seasonal components.

### Quantile Regression Forest

Random forests are collections of bagged regression trees [14]. Upon sampling a random set of predictors, each tree generates one prediction for the next period’s count of ILI positive visits. A random forest forecast consists of the average of the predictions of all trees.

We implemented an extension to random forests, called quantile regression forest (QRF), using the *quantregForest* R package [15]. QRF generalizes the usual random forest prediction by estimating conditional quantiles, useful when constructing prediction intervals. The predictors included in the QRF model were: lags of weekly ILI counts; current epi-week^3^; lags of weekly minimum and maximum temperature; lags of weekly flu immunization rate; and city population. Other predictors were considered but were discarded by recursive feature elimination using the rfe method of the R *caret* package [16]. These included weekly counts of diagnosed cases of influenza A H1N1 and H3, influenza B, rhino/enterovirus, and respiratory syncytial virus.^4^ Information on patient age, sex, and location within city were also excluded since they rendered the model computationally expensive and yielded no performance gains.^5^

Our implemented QRF used *n* = 2000 decision trees, sampling *m* = 4 of the available variables each time. The parameter *m* was chosen using the conventional heuristic 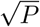, where *P* represents the number of predictors. The QRF prediction of ILI positive visits *ℓ* weeks from week *t*, 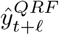, is given by:

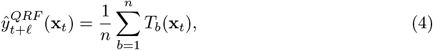

where each *T_b_* is one of the n decision trees resulting from a bootstrapped sample of the training data with m randomly selected predictors and **x**_*t*_ are the values of the model predictors at week t. The prediction interval of the QRF stems from conditional quantiles computed by the *quantregForest* method in R. [17].

### Linear Regression

We fitted a standard linear regression (LR) model with the following predictor variables: current week’s ILI positive visit count; weekly-averaged minimum and maximum temperatures; city population; immunization rate; year trend; epi-week fixed effects; slope of the ILI curve at current period; and, a categorial variable counting upward movements in weekly ILI positive visits over the preceding three weeks. The ILI slope and count of upward movements helped on detecting sharp increases in recent ILI positive visits and improved the prediction near the peak. The model’s prediction of ILI positive visits *ℓ* weeks from week *t* is given by:

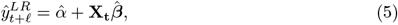

where **X**_t_ is a row vector of the values for the predictors at week *t*, 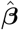 is a column vector with the estimated marginal effect of each predictor on the ℓ weeks ahead count of ILI positive visits and 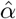 is the intercept estimate. The prediction interval is 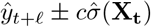 where 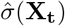 is the standard error of the prediction given observed values **X**_t_ and *c* once again comes from the amount of coverage for the prediction interval under the assumption that errors are normally distributed.

### Stacked Ensemble

We computed a weighted-average prediction 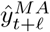 based on the predictions of *M* contributing models {*EB,QRF,ARIMA, LR*} described above:

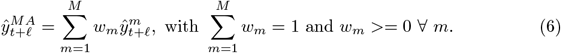

We used stacking, a data-driven approach, to find a set of weights *w_m_* for combining the predictions of individual models in a manner that minimized prediction error in a held out data set. Specifically, we found the weights:

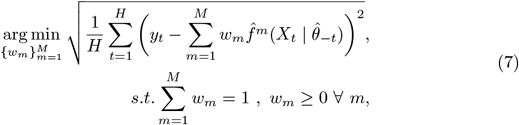

where *H* is the size of the hold-out data and 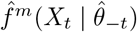 is the prediction at period *t* of model *m* trained without data from the flu season encompassing period *t*. In practice, the algorithm for deriving the stacked ensemble weights is as follows:

1. Train each individual model to the dataset holding out all weeks comprising an flu season;
2. Obtain fitted values for the weeks in the hold-out data;
3. Repeat steps 1 and 2 using every other flu season as hold-out data;
4. Compute the weights using eq 7 with the *H* weeks of predictions obtained in steps above.

The rationale for ensembling is its potential to reduce prediction error by reducing prediction variance and, in some cases, bias [19]. The benefits of ensembling are typically smaller as the predictions of individual models become more positively correlated. Furthermore, as shown by Claeskens *et al* [20], estimation of averaging weights introduces additional randomness to the ensembled prediction. As such, we also show results for a “naïve” ensemble using equal weights on each individual model, i.e. the case where 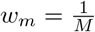 for each of the *M* models contributing to the ensembled prediction.

Deriving the sampling distribution of 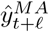 is nontrivial as the ensemble mixes the distributions of each contributing model and some individual models are non-parametric to begin with. The literature on model averaged prediction intervals is scarce, and even in simpler contexts with parametric models the constructed intervals perform poorly in terms of coverage rate on validation exercises [21]. We report a weighted average of quantiles of the distribution of each individual model prediction as the ensembles’ own distributions. In our analysis, we show the coverage rate of this back-of-the-envelope ensemble prediction interval, as well as the coverage rate of intervals constructed from each individual model.

### Analysis

Each model was trained on a partition of the available data and subsequently evaluated against held out data. The model evaluation used a leave-one-out approach: each flu season was held out once and used as a test set, with the model being trained on the remaining seasons. Note that the test set was removed before executing the algorithm for the ensemble methods to ensure that test data did not contribute to the construction of the ensemble weights. We considered four commonly used metrics for comparing model performance: mean absolute error (MAE); root mean square error (RMSE); mean absolute percentage error (MAPE); and log scoring. These are described below.

### Mean Absolute Error & Root Mean Square Error

Our forecasting targets were the ℓ ∈ {1, 2, 3, 4} weeks ahead ILI positive visit counts. We summarized model performance for each target using standard measures of prediction error: MAE and RMSE. These are given by:

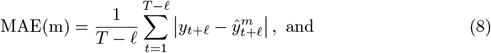

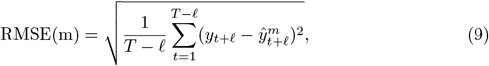

where *y*_*t*+ℓ_ is the realized weekly ILI positive count at period *t* + ℓ and 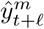 is model *m*’s generated ℓ weeks ahead prediction made at period *t*. MAE and RMSE express an *average* prediction error over the *T* – ℓ observations in the sample. Both metrics are increasing with the average error, meaning the models with best accuracy in the test set are the ones with the lowest MAE and RMSE. Squaring of the error implies the RMSE imposes a greater penalty for larger deviations between predicted and observed values relative to the MAE.

### Mean Absolute Percentage Error

For the sake of interpretability, we also present each model’s MAPE, which expresses the average percentage deviation between forecast and realized outcomes. The MAPE formula is given by:

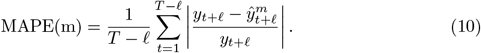

MAPE is known to produce infinite or undefined values when the denominator of any summation term in eq 10 approaches zero [22]. This disadvantage did not apply to our analysis since we had a minimum of 52 weekly ILI visits to the ED during the sample period.

### Log Score

The last performance metric in our analysis is a variant of the log score measure used to rank participants on the United States Centers for Disease Control (US CDC) flu forecasting challenge [23]. The log score of a prediction measures how much probability our model assigns to an “acceptable” prediction range. We defined a prediction as acceptable if it fell within ± 25 visits of the realized weekly ILI positive weekly count.

Since we compare a mix of parametric and non-parametric models, we approximate the probability assigned to the acceptable range by the count of centiles of the forecast inside the range. That is, the log score of a ℓ weeks ahead prediction from model *m* at week *t* is computed as follows:

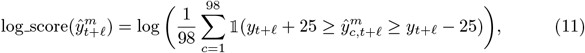

where 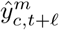 is the *c* centile of the (probabilistic) forecast generated by model m. In the event the observed ILI visits fell below the first centile or above the 99th centile, we assigned the value —5 to the log score, which is slightly lower than log(0.01) ≈ −4.6.

### Descriptive Statistics

Table 1 summarizes key variables used in the analysis. The high visit volume ED at ACH received on average 208 ILI positive visits per week during the sample period^6^. This average increased almost 60% to 332 ILI positive visit counts during high visit volume weeks – i.e. those on the top 25% of ILI visit counts – within each season. This increased pattern was also visible for lab confirmed cases of different virus strains, except for rhino/enteroviruses which varied less predictably throughout the flu season. The last column of table 1 shows the *t* statistic for the difference in means between high and low visit volume weeks – i.e. weeks on the bottom 75% of ILI visit counts – within each season. In addition to the mechanical increase in ILI during high volume weeks, we see as expected lower temperatures during high volume weeks, but little evidence of differences in patient demographic characteristics and flu immunization rates by visit volume.

**Table 1.**
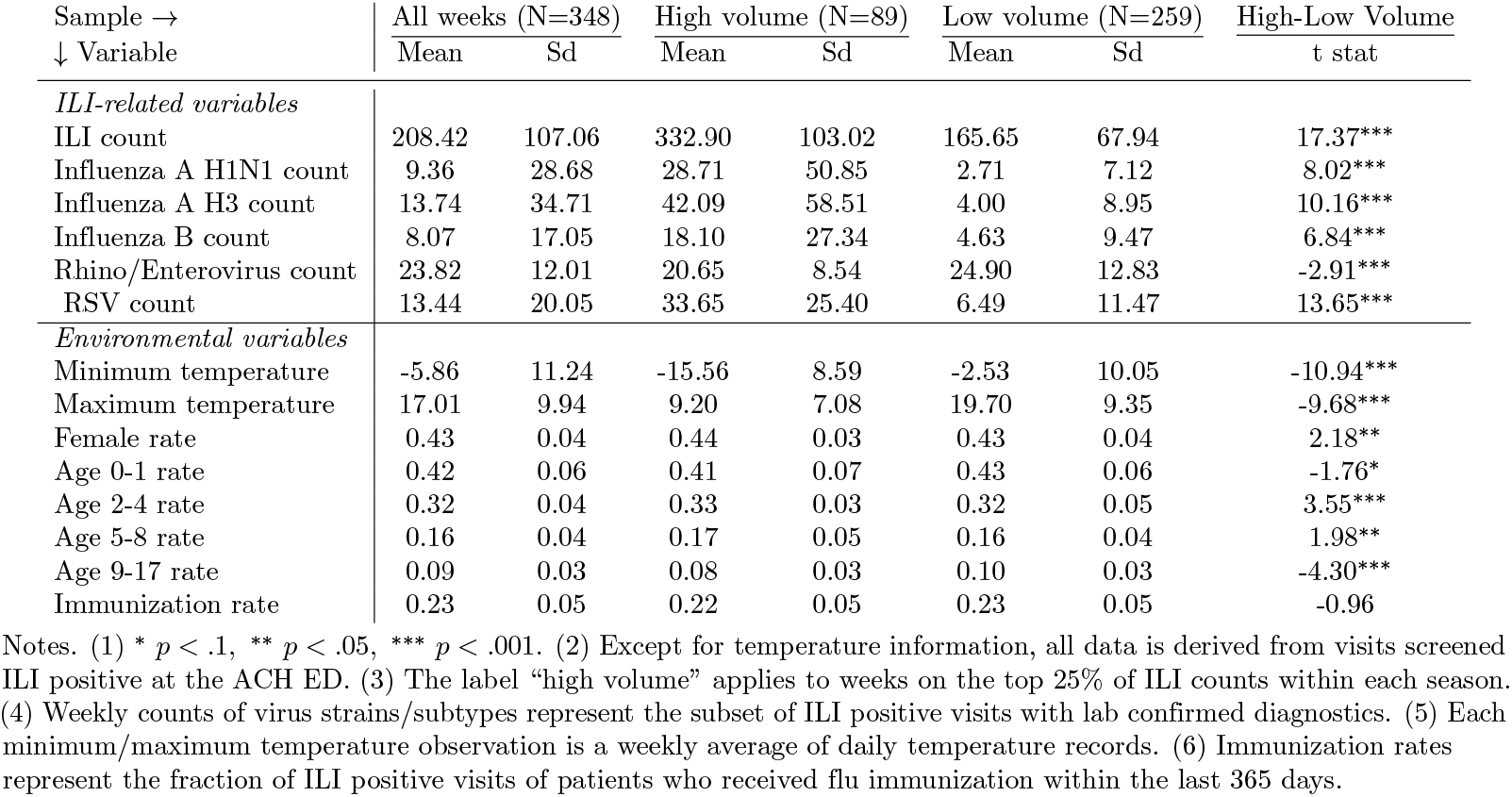
Descriptive statistics.

## Results

### Overall comparison

We compared each model’s performance metrics over all prediction targets. Fig 1 and Fig 2 show the MAE and RMSE of all models for the 1 to 4 weeks ahead predictions. These figures illustrate how predictions worsen as we look further into the future. Both MAE and RMSE yield the same ordering of model performance, except for a higher RMSE on the 3 and 4 weeks forecast of the EB model. This suggests EB predictions are farther from the truth on the longer forecast horizons. Fig 1 and Fig 2 also highlight the fact that predictions which draw information solely on the trajectory of ILI (EB and ARIMA forecasts) perform considerably worse than predictions modeling the relationship between environmental factors such as temperature and immunization rates. Fig 3 shows the best performing model’s (stacked ensemble) 1 week ahead predictions deviate from the observed ILI count by an average of 12% for 1 week ahead and 19% for 4 weeks ahead predictions, in absolute terms.

**Fig 1.**
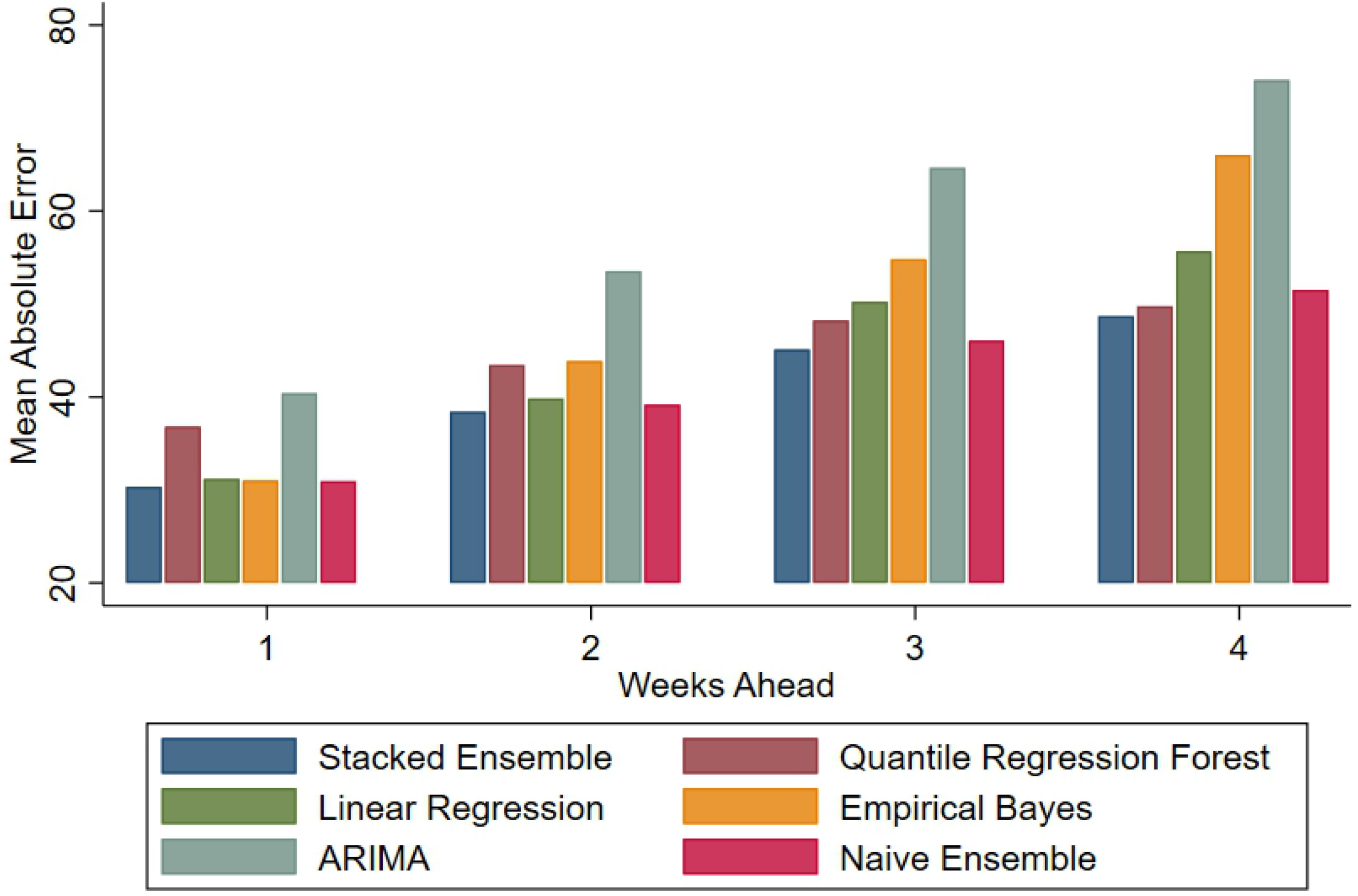
MAE comparison for test seasons 2012-2018.

**Fig 2.**
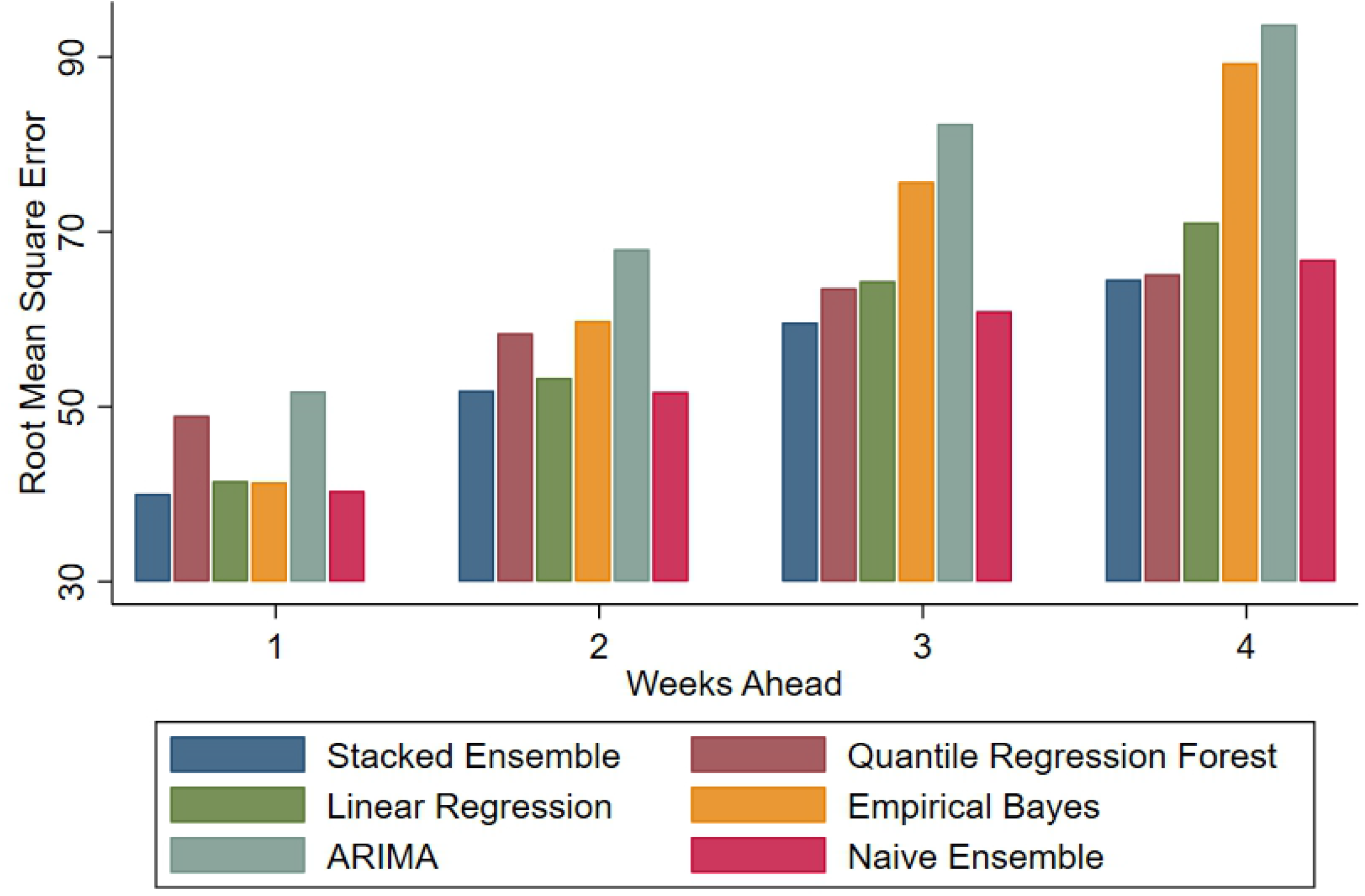
RMSE comparison for test seasons 2012-2018.

**Fig 3.**
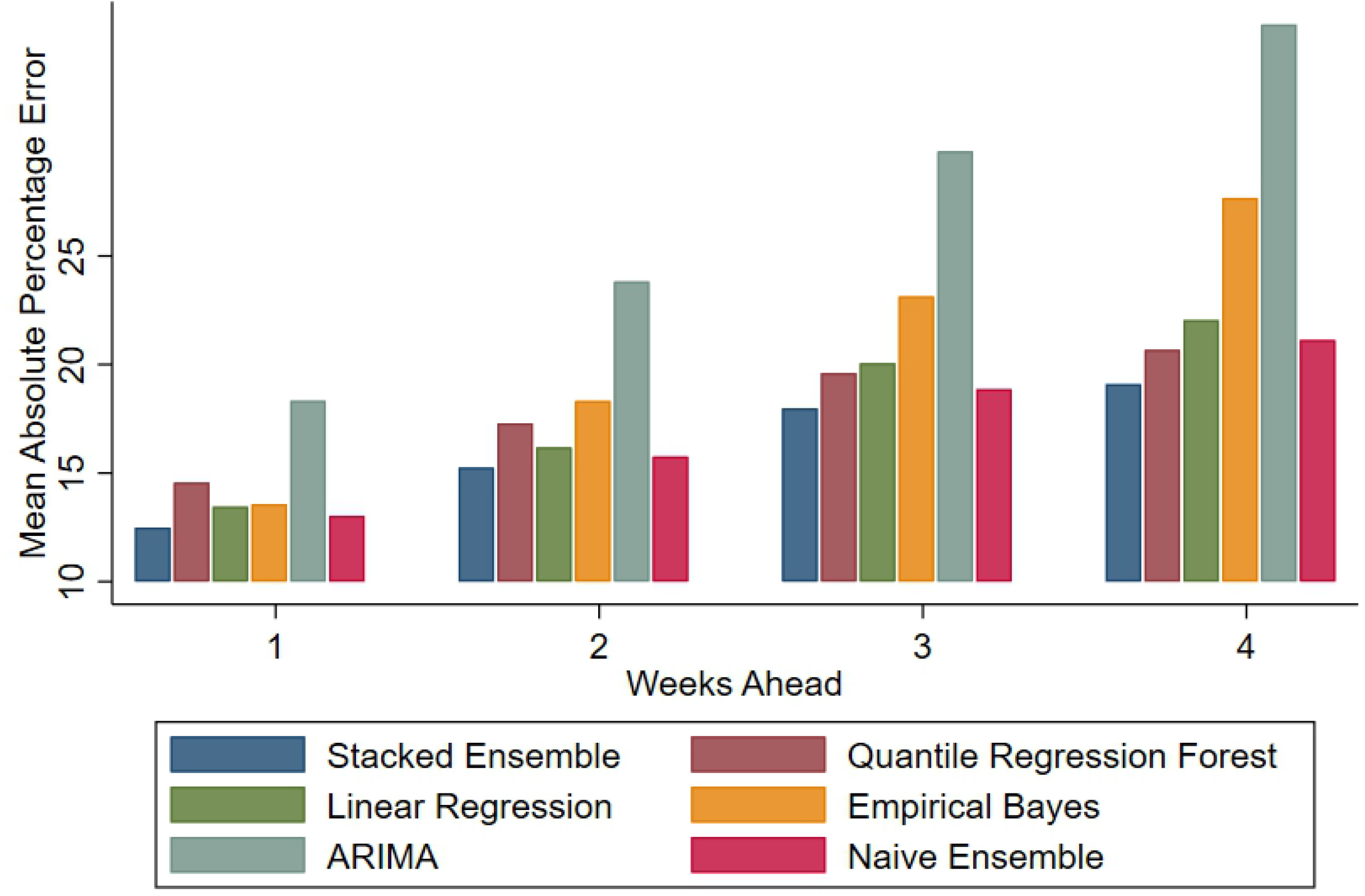
MAPE comparison for test seasons 2012-2018.

This overall comparison of models suggests the stacked ensemble was competitive across all performance metrics and all prediction targets. Predicted counts generated by this approach deviated, on average, by 30 and 50 weekly ILI visits in the nearest 1 week and farthest 4 weeks ahead targets, respectively. In fact, forecasts generated by combining predictions from various models are gaining popularity as they tend to be on average more accurate than the individual models they combine.

#### Flu season breakdowns

The results presented in the overall comparison subsection paint a general picture about each model’s ability to predict future ILI weekly counts. Table 2 provides a more granular perspective by examining model performance separately for each season. Table columns alternately report the models’ mean log score for each season. The stacked ensemble outperforms individual models for the 1 and 2 weeks ahead forecast, falling slightly behind the QRF on the more distant forecasts. Table 3 reports for each model and flu season the percentage of predictions that fall within 20% of the observed weekly counts on short-term, 1 week ahead, forecasts. Each column represents a prediction target for a specific flu season. The stacked ensemble method produced the highest percentage of predictions within 20% in more seasons than any other model, also ranking highest on the last two columns aggregating predictions over all seasons.

**Table 2.** Mean log score of each model, by flu season and forecast horizon.

| Flu Season →<br>↓ Model | <u>2012-13</u> | <u>2013-14</u> | <u>2014-15</u> | <u>2015-16</u> | <u>2016-17</u> | <u>2017-18</u> | <u>2018-19</u> | <u>All Seasons</u> |
| --- | --- | --- | --- | --- | --- | --- | --- | --- |
| *1 week ahead forecast* |  |  |  |  |  |  |  |  |
| Empirical Bayes | -1.13 | -1.01 | -1.31 | -1.54 | -1.11 | -1.12 | -2.47 | -1.27 |
| ARIMA | -2.10 | -1.81 | -1.77 | -1.59 | -1.66 | -1.46 | -4.42 | -1.88 |
| Quantile Regression Forest | -1.10 | -0.94 | -1.14 | -1.11 | -1.21 | -1.26 | -2.87 | -1.22 |
| Linear Regression | -1.17 | -0.95 | -1.18 | -1.53 | -1.08 | -1.06 | -2.63 | -1.24 |
| Stacked Ensemble | **<u>-0.96</u>** | **<u>-0.89</u>** | -1.06 | **<u>-1.10</u>** | **<u>-1.02</u>** | -0.99 | **<u>-2.24</u>** | **<u>-1.07</u>** |
| Naive Ensemble | -1.02 | -0.92 | **<u>-1.05</u>** | **<u>-1.10</u>** | -1.03 | **<u>-0.98</u>** | -2.58 | -1.10 |
| *2 weeks ahead forecast* |  |  |  |  |  |  |  |  |
| Empirical Bayes | -1.39 | -1.18 | -1.39 | -1.58 | -1.52 | -1.37 | **<u>-2.77</u>** | -1.48 |
| ARIMA | -2.66 | -2.62 | -1.53 | -1.97 | -2.27 | -1.56 | -5.84 | -2.31 |
| Quantile Regression Forest | -1.20 | **<u>-1.10</u>** | -1.27 | **<u>-1.26</u>** | -1.32 | -1.41 | -3.51 | -1.38 |
| Linear Regression | -1.24 | -1.20 | -1.41 | -1.63 | -1.43 | -1.20 | -3.83 | -1.49 |
| Stacked Ensemble | **<u>-1.15</u>** | **<u>-1.10</u>** | -1.18 | -1.26 | -1.22 | -1.12 | -3.35 | **<u>-1.29</u>** |
| Naive Ensemble | -1.21 | -1.18 | **<u>-1.14</u>** | **<u>-1.21</u>** | **<u>-1.21</u>** | **<u>-1.08</u>** | -3.39 | **<u>-1.29</u>** |
| *3 weeks ahead forecast* |  |  |  |  |  |  |  |  |
| Empirical Bayes | -1.60 | -1.42 | -1.95 | -1.83 | -1.72 | -1.35 | -3.42 | -1.74 |
| ARIMA | -3.28 | -3.26 | -1.93 | -2.32 | -2.86 | -1.57 | -5.71 | -2.71 |
| Quantile Regression Forest | **<u>-1.19</u>** | -1.14 | **<u>-1.34</u>** | **<u>-1.36</u>** | **<u>-1.32</u>** | -1.47 | **<u>-3.19</u>** | **<u>-1.41</u>** |
| Linear Regression | -1.41 | -1.38 | -1.38 | -1.80 | -1.67 | -1.33 | -5.07 | -1.69 |
| Stacked Ensemble | -1.24 | **<u>-1.11</u>** | -1.38 | -1.46 | -1.38 | -1.26 | -3.57 | -1.43 |
| Naive Ensemble | -1.37 | -1.32 | -1.27 | -1.39 | -1.39 | **<u>-1.19</u>** | -4.00 | -1.47 |
| *4 weeks ahead forecast* |  |  |  |  |  |  |  |  |
| Empirical Bayes | -1.78 | -1.58 | -1.84 | -1.97 | -1.84 | -1.52 | -3.62 | -1.86 |
| ARIMA | -3.95 | -3.82 | -1.94 | -2.55 | -3.05 | -1.81 | -6.66 | -3.06 |
| Quantile Regression Forest | **<u>-1.24</u>** | **<u>-1.14</u>** | -1.29 | **<u>-1.44</u>** | **<u>-1.34</u>** | -1.41 | **<u>-3.02</u>** | **<u>-1.41</u>** |
| Linear Regression | -1.54 | -1.45 | -1.43 | -1.77 | -1.78 | -1.43 | -5.04 | -1.76 |
| Stacked Ensemble | -1.40 | -1.21 | **<u>-1.25</u>** | -1.53 | -1.53 | **<u>-1.27</u>** | -3.59 | -1.49 |
| Naive Ensemble | -1.45 | -1.38 | -1.27 | -1.51 | -1.53 | **<u>-1.27</u>** | -4.18 | -1.55 |

Notes. Due to data availability, the 2018 season uses only 14 weeks of data, starting from epiweek 37 or approximately mid September.

**Table 3.** Percentage of 1 week ahead predicted ILI visits within 20% of observed visits, by flu season.

| Flu Season →<br>↓ Model | <u>2012-13</u> | <u>2013-14</u> | <u>2014-15</u> | <u>2015-16</u> | <u>2016-17</u> | <u>2017-18</u> | <u>2018-19</u> | <u>All Seasons</u> |
| --- | --- | --- | --- | --- | --- | --- | --- | --- |
| *All weeks in sample* |  |  |  |  |  |  |  |  |
| Empirical Bayes | 76.7 | 76.7 | 69.8 | 67.4 | 79.1 | 88.4 | **80.0** | 76.6 |
| ARIMA | 48.8 | 58.1 | 74.4 | **79.1** | 76.7 | 86.0 | 66.7 | 70.3 |
| Quantile Regression Forest | 60.5 | **88.4** | 69.8 | **79.1** | **86.0** | 90.7 | 60.0 | 78.0 |
| Linear Regression | 67.4 | 83.7 | 74.4 | 74.4 | 81.4 | 93.0 | 66.7 | 78.4 |
| Stacked Ensemble | **79.1** | 83.7 | **79.1** | 72.1 | **86.0** | **95.3** | **80.0** | **82.4** |
| Naive Ensemble | 72.1 | 83.7 | 74.4 | 69.8 | 83.7 | **95.3** | 73.3 | 79.5 |
| *High volume weeks* |  |  |  |  |  |  |  |  |
| Empirical Bayes | 63.6 | 83.3 | **90.9** | 81.8 | 81.8 | 90.9 | **100.0** | 83.1 |
| ARIMA | 54.5 | 58.3 | 81.8 | 81.8 | 72.7 | 90.9 | 75.0 | 73.2 |
| Quantile Regression Forest | 72.7 | 83.3 | **90.9** | **90.9** | 81.8 | **100.0** | 25.0 | 83.1 |
| Linear Regression | 63.6 | **100.0** | 72.7 | 81.8 | **90.9** | **100.0** | 75.0 | 84.5 |
| Stacked Ensemble | **81.8** | 83.3 | **90.9** | **90.9** | **90.9** | **100.0** | 75.0 | **88.7** |
| Naive Ensemble | **81.8** | 91.7 | **90.9** | 81.8 | 81.8 | **100.0** | 75.0 | 87.3 |

Notes. (1) Due to data availability, the 2018 season uses only 14 weeks of data, starting from epiweek 37 or approximately mid September.

### Prediction interval coverage rate

As discussed in Section 2.2, each model uses a different approach when characterizing the uncertainty of the predictions. In sum, constructing prediction intervals for the ARIMA and LR models is straightforward and follows from estimates of the standard error of the prediction. Meanwhile the EB estimates have a prediction interval computed from quantiles of the posterior distribution and the RF prediction interval stems from quantiles of the output from individual regression trees. Lastly the interval around the stacked ensemble estimate (as well as naive ensemble) is a back-of-the-envelope calculation from weighting the quantiles of the predictions from individual models.

We assessed the coverage of each model’s prediction interval on the complete test seasons from 2012-13 to 2017-18. Fig 4 shows, for each model and flu season, the 1 week ahead prediction against the realized ILI visit counts in that particular week. The shaded region corresponds to a 90% prediction interval generated by the model and the coverage rate refers to the percentage of observed weekly ILI visits that falls inside the model’s prediction interval. The figures show ARIMA prediction intervals had the lowest coverage, potentially since uncertainty associated with the STL decomposition was ignored. Except for ARIMA and the EB model, the coverage rates of models were slightly higher than 90% and as such the intervals on our most competitive models can be viewed as conservative estimates.

**Fig 4.**
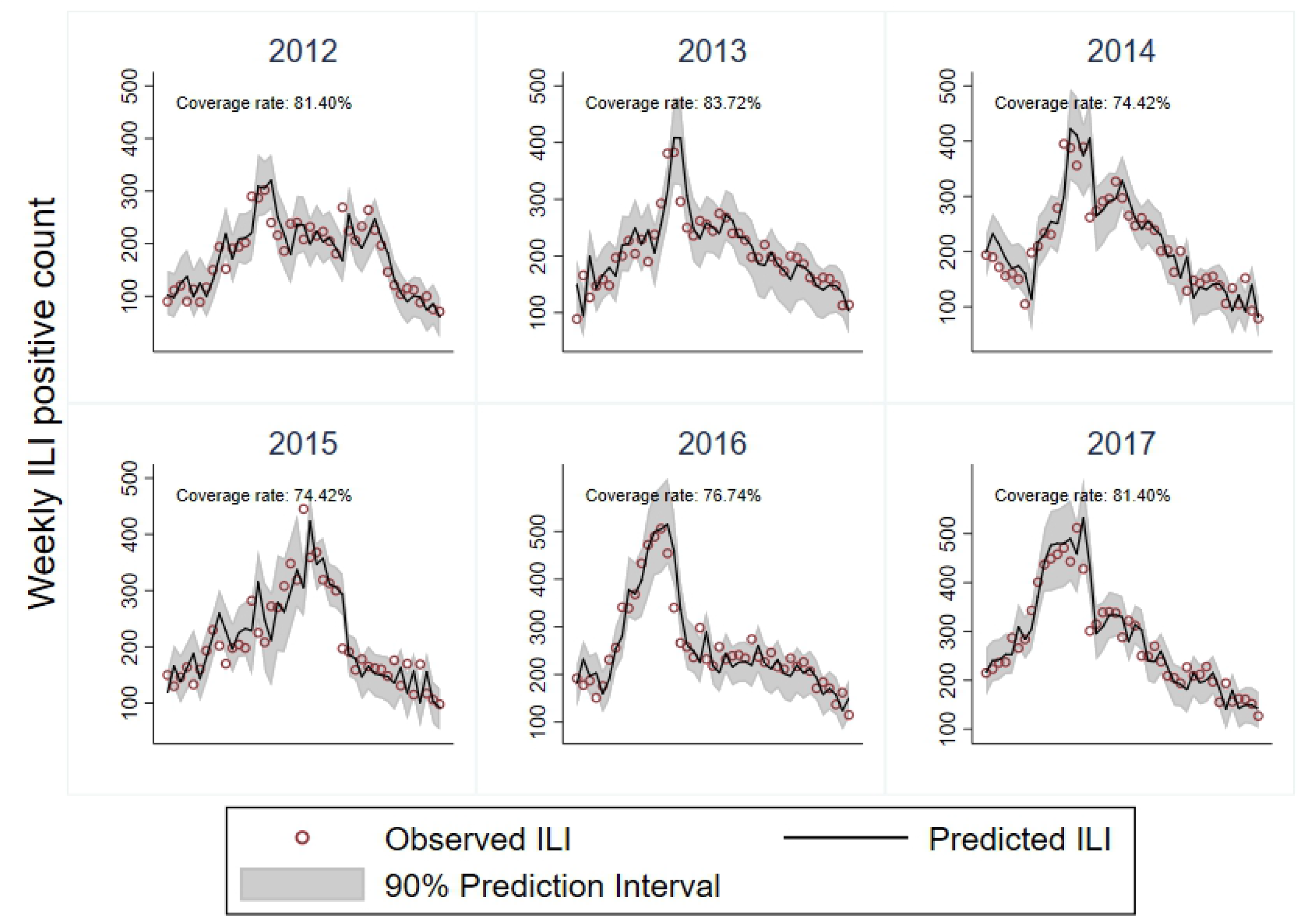
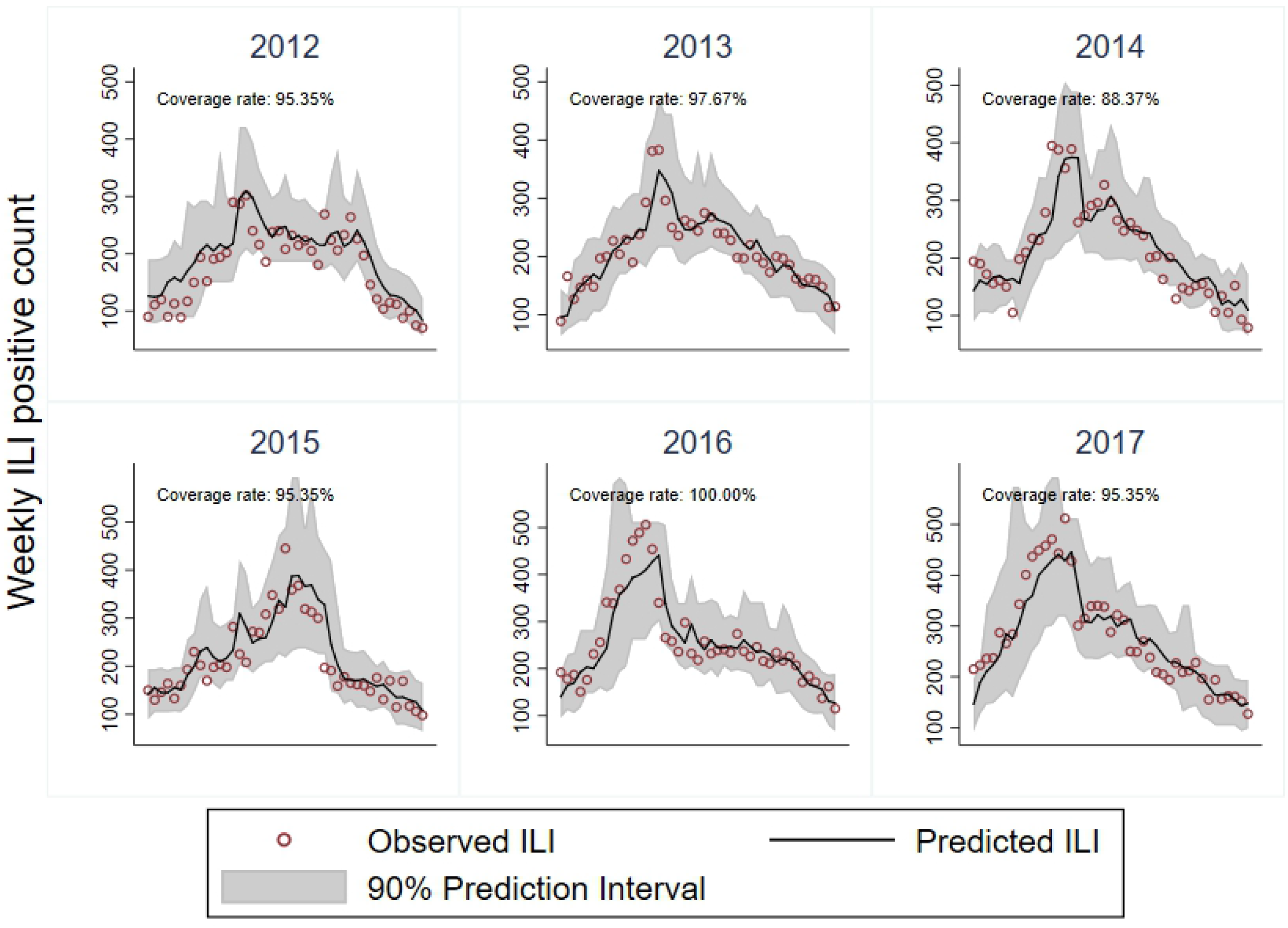
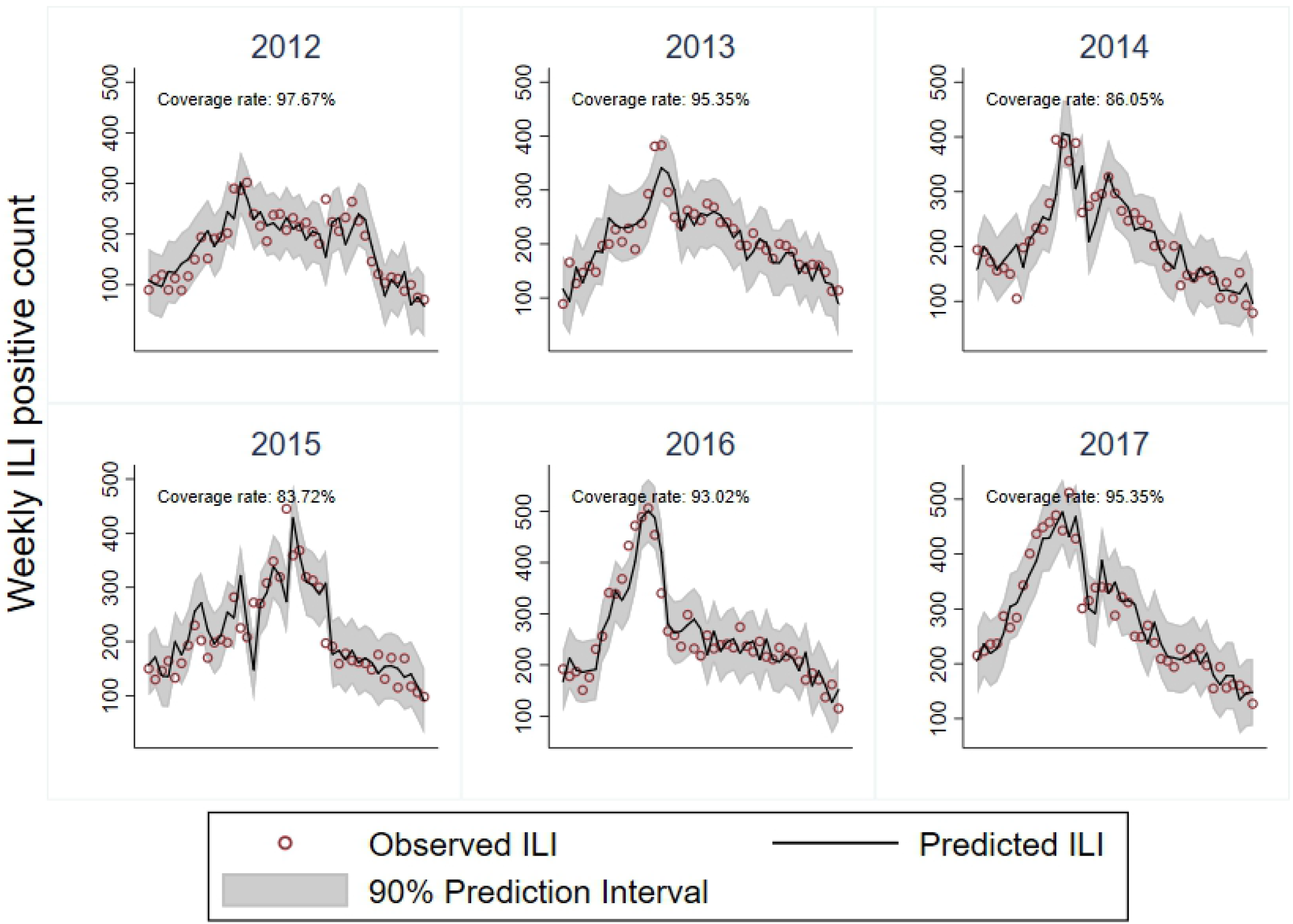
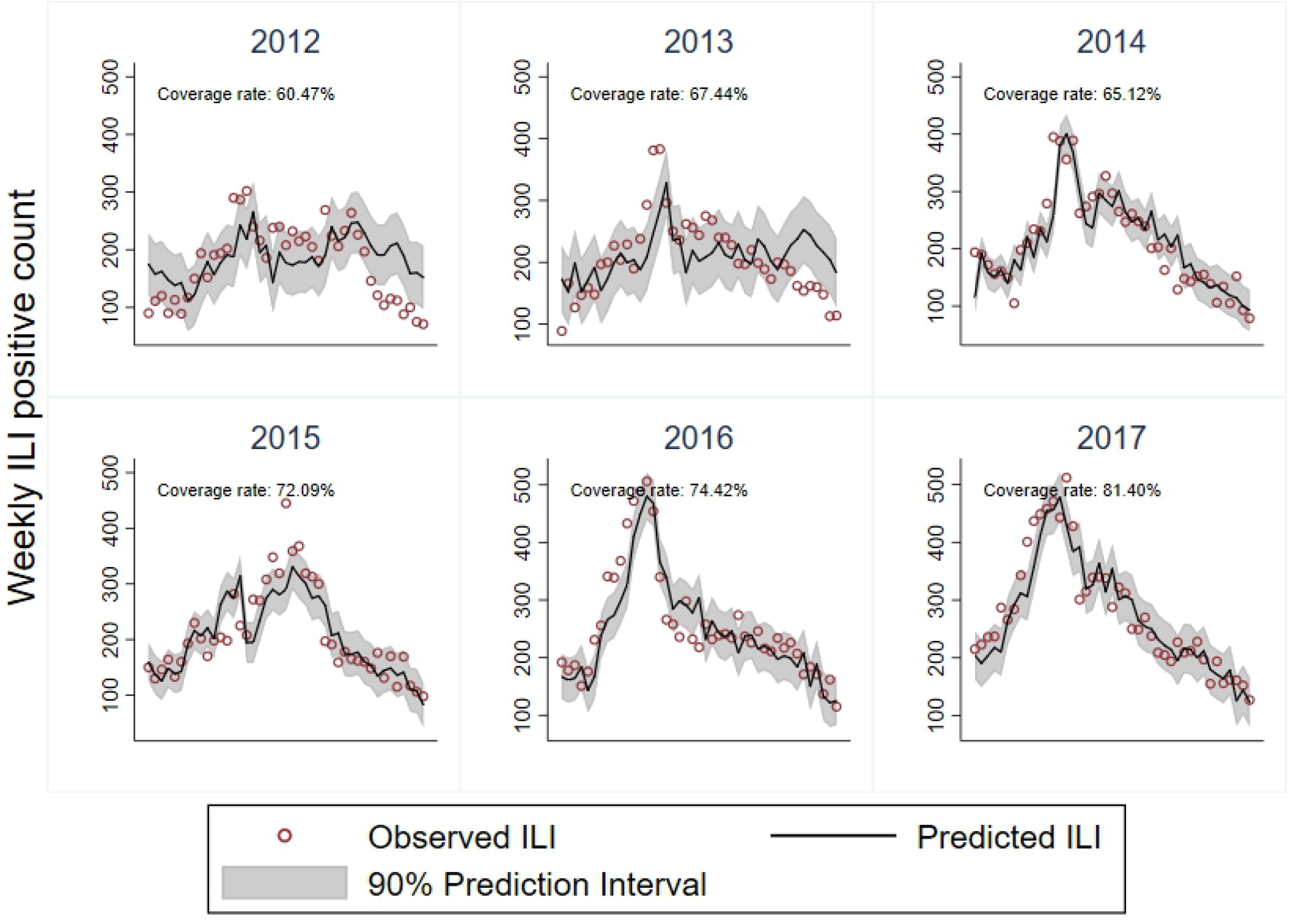
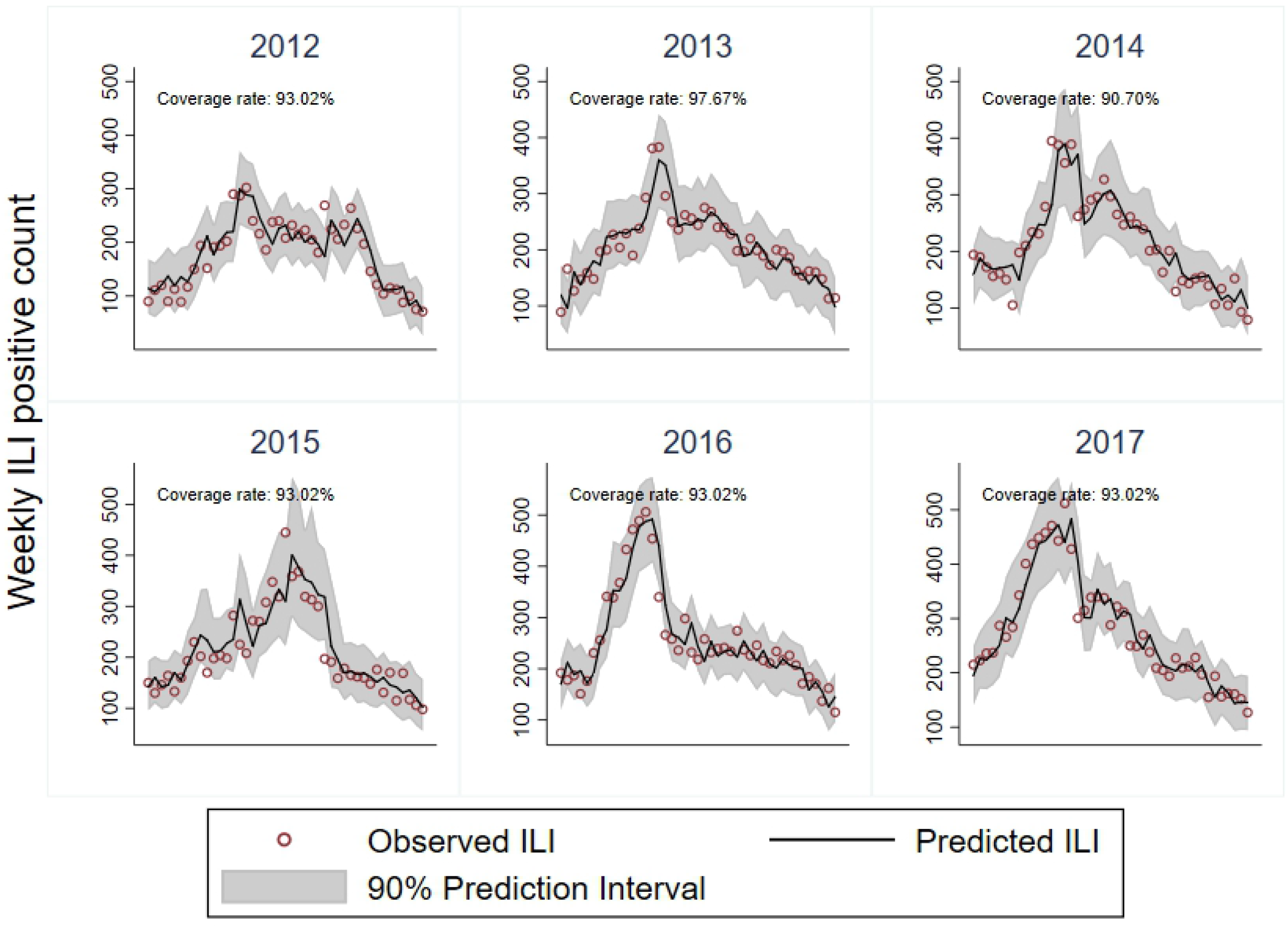
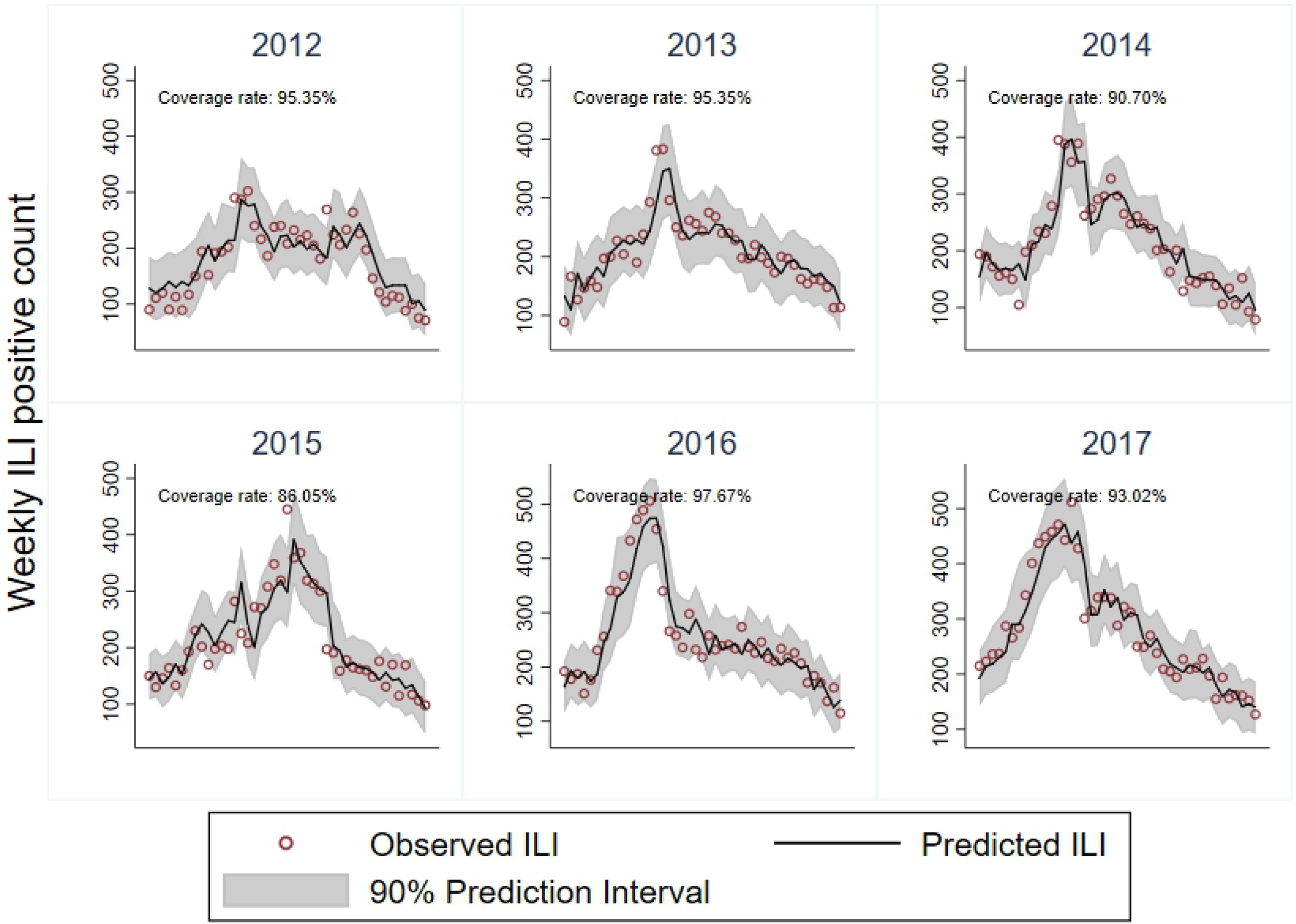
Predicted vs Observed ILI visits for 1 week forecast. A: Empirical Bayes. B: Quantile Regression Forest. C: Linear Regression. D: ARIMA. E: Stacked Ensemble. F: Naive Ensemble.

### Alternative data set

We evaluated whether our findings were robust to using data from the lower visit volume ED at FMC hospital, located in the vicinity of ACH in Calgary, Alberta. S2 Fig shows the RMSE of each model when trained and evaluated on data from the FMC ED. Results were qualitatively consistent with those in the main dataset, with the stacked ensemble still outperforming the individual models. In addition to lower visit volumes, the FMC ED caters to an adult population, in contrast to the children’s hospital used in the main analysis.

## Discussion

We tested the performance of multiple models for predicting ILI visits at the ED level. Our findings promote stacked ensembling – a data-driven model averaging method – as a viable approach for short term forecasts of weekly ILI visits. We show, through various exercises comparing model performance, that our stacked ensemble prediction leverages the strength of individual models and therefore provides a robust estimate on both short-term (1 week) and longer term (4 weeks) forecast horizons.

Previous research on forecasting ILI have discussed potential improvements of incorporating demographic and spatial information into the statistical models. We found no evidence of such improvements in our research design, possibly due to small sample size and the potential for overfitting when including additional predictors. Instead, our preferred model required only weekly counts of ILI visits to the ED, the rate of ILI patients who are immunized in the current season, and temperature and population data which are typically available in urban areas. In addition, the tested models are easily implementable with free software for statistical computing.

We found that predictions from our stacked ensemble outperformed those of individual models in two distinct EDs located in Calgary, Alberta. Future work should include a larger sample of hospitals in the province and an attempt at modelling cross effects of ILI visits in an ED on the prediction of ILI for neighboring EDs. This could potentially improve predictive power at lower visit volume EDs where outbreak detection could be of relevance.

## Supporting information

**S1 Fig. RMSE comparison for Quantile Regression Forests with added features.** Notes. (1) Data from weekly ILI visits at the ACH. (2) The Baseline QRF is described in section 2.2; the other QRF specifications build on the Baseline. E.g., Sex means we disaggregate weekly ILI counts by male and female patients. (3) Calculation of RMSE uses 42 weeks on each test season of the ACH data, starting from epiweek 37 or approximately mid September. (4) Due to data availability, the 2018 season uses only 14 weeks of data, also starting from epiweek 37.

**S2 Fig. RMSE comparison for test seasons 2012-2018 (FMC hospital).**

## Acknowledgments

We thank members of Public Health Surveillance and Infrastructure at Alberta Health Services. Hussain Usman and Adrienne Macdonald provided helpful input and feedback during the model development stage. We also thank Sherry Trithart and Shaun Malo for data provision from ARTSSN.

## Footnotes

1 The laboratory data reports diagnostic results for Adenovirus; Coronavirus; HMPV; Influenza A H1N1; Influenza A H3; influenza A (untyped); influenza B; parainfluenza; rhino/enterovirus; and, RSV.

2 A prediction interval was also constructed from percentiles of the *N* weighted samples.

3 Epi-weeks follow the US CDC version of an epidemiological week. It adopts the same rules as ISO weeks, but starts on Sunday. In our analysis, epi-weeks were constructed using the epiweek method of the *lubridate* R package [18].

4 These five represent the most frequent strain types in the data. We avoided including too many covariates to keep the model parsimonious and mitigate the risk of overfitting.

5 To illustrate, the S1 Fig shows the effect of adding patient sex, age, sex and age, location within city, or strain types on the performance of the “baseline” QRF. The RMSE performance metric in this figure is described in the analysis section of the paper.

6 In contrast, the lower visit volume ED at FMC received on average 31 ILI visits per week during the same period.

